# Improving brain age estimates with deep learning leads to identification of novel genetic factors associated with brain aging

**DOI:** 10.1101/2020.07.08.192617

**Authors:** Kaida Ning, Ben A. Duffy, Meredith Franklin, Will Matloff, Lu Zhao, Nibal Arzouni, Fengzhu Sun, Arthur W. Toga

**Affiliations:** USC Stevens Neuroimaging and Informatics Institute, Keck School of Medicine of University of Southern California, Los Angeles, California 90033, USA; Molecular and Computational Biology Program, University of Southern California, Los Angeles, CA 90089, USA; Department of Preventive Medicine, University of Southern California, Los Angeles, CA 90032, United States; Neuroscience Graduate Program, University of Southern California, Los Angeles, CA 90089, USA

**Author notes:** These authors contributed equally to this work (Kaida Ning, Ben A. Duffy). Corresponding author at: Arthur W. Toga, USC Stevens Neuroimaging and Informatics Institute, Keck School of Medicine of University of Southern California, 2025 Zonal Ave., Los Angeles, California 90033, USA.

## Abstract

Brain aging trajectories among those of the same chronological age can vary significantly. Statistical models have been created for estimating the apparent age of the brain, or predicted brain age, with imaging data. Recently, convolutional neural networks (CNNs) have shown the potential to more accurately predict brain age. We trained a CNN on 16,998 UK Biobank subjects, and in validation tests found that it was more accurate than a regression model for predicting brain age. A genome-wide association study was conducted on CNN-derived predicted brain age whereby we identified single nucleotide polymorphisms from four independent loci significantly associated with brain aging. One locus has been previously reported to be associated with brain aging. The three other loci were novel. Our results suggest that a more accurate brain age prediction enables the discovery of novel genetic associations, which may be valuable for identifying other lifestyle factors associated with brain aging.

## 1 Introduction

Brain structural changes that occur during the brain aging process vary between individuals. A long-standing question is how genetic and environmental factors contribute to this variation.

Recently, statistical models have been developed to capture structural changes and assess the apparent age of the brain based on imaging data (*1-3*). The biomarker predicted from these models is referred to as the predicted brain age (PBA). By comparing PBA and chronological age, researchers can quantify whether a subject’s brain appears older or younger than expected, which is reflective of accelerated or decelerated aging of the brain. This has allowed identification of genetic factors, lifestyle factors, and diseases that are associated with accelerated brain aging (*4-7*).

Genetic factors associated with brain aging are important to identify because they could allow clinicians to target brain aging at an early stage and potentially facilitate effective interventions. Two recent studies investigated the association between single nucleotide polymorphisms (SNPs) and brain aging using UK Biobank data. Jonsson et al. used a CNN model for predicting brain age where the predictor was the whole 3D MRI scan (*4*), while Ning et al. used a linear regression model for predicting brain age, where predictors were brain morphometric measurements derived from MRI scans (*5*). Both studies reported mean absolute error (MAE) between PBA and the true chronological age around 3.5 years and found an association between a chromosome 17 locus and brain aging. That locus includes the MAPT gene, mutations in which are associated with dementia and Parkinson’s disease (*8*).

Recent advances in training CNNs, which are deep learning models that learn features in images with convolution operations without prior knowledge of what these features are, have allowed researchers to accomplish disease and phenotype classification and prediction with high accuracy (*9-11*). Recently, Langner et al. trained a CNN for predicting age of the body based on whole-body MRI of about 20,000 subjects. The MAE between predicted ‘body age’ and true chronological age reached 2.5 years (*11*). In comparison, in a recent study on the association between genetic variants and brain aging where PBA was obtained using a CNN model, the model was trained with fewer than 2,000 subjects and had MAE around 3.5 years (*4*).

In our study, we used a much larger sample of brain imaging data of 16,998 UK Biobank subjects for training a CNN to obtain PBA. Following this, we calculated relative brain age (RBA), a metric describing a subject’s PBA relative to peers (*5*). We then conducted a genome-wide association study on this CNN-derived RBA to identify SNPs significantly associated with RBA. We further compared the SNPs associated with the CNN-derived RBA to those found using a linear regression. Our goal was to leverage a large sample to investigate whether the CNN-derived RBA metric is more accurate than linear regression and to test whether it enables the discovery of a greater number of genetic variants associated with brain aging. If a CNN-derived RBA metric is both more accurate and reveals more genetic variants associated with brain aging, it would suggest that CNN could also be used to more effectively investigate the association of lifestyle and disease factors with brain aging.

## 2 Results

### 2.1 Predicted brain age from a CNN model

In the five evaluation sets, the median MAE between CNN-derived PBA and chronological age was 2.7 years (ranging from 2.5 years to 3.1 years). We observed that the difference between PBA and chronological age (i.e., PBA-CA) was negatively associated with chronological age. Older subjects had negative PBA-CA, while younger subjects had positive PBA-CA (Supplementary Fig. 1). Therefore, after obtaining PBA, we adjusted for this association by calculating the RBA metric, which is independent of age, and assessed the association between RBA and genetic variants.

### 2.2 Genetic factors associated with brain aging

In total, 80 SNPs from four independent loci show significant associations with RBA derived from CNN (association p-value < 5E-8). Fig. 1 shows a Manhattan plot of the associations between SNPs and RBA across the genome. Supplementary Table 1 lists the SNP-level association p-values (DOI:10.5281/zenodo.3786826). Three SNPs from a locus on chromosome 4 showed significant association with RBA, with the leading SNP rs337638 having an association p-value of 3E-16. These SNPs are located in a non-coding region with the nearest protein coding gene being KLF3 (Fig. 2). Multiple SNPs significantly associated with RBA are located in a chromosome 17 locus that has been previously linked to brain aging (*4, 5*). This locus includes genes such as NSF, MAPT, and WNT3 (Figure3). In addition, SNP rs2677110 located within the gene RUNX2 on chromosome 6 and SNP rs12775208 located within the gene NKX6-2 on chromosome 10 also showed significant association with RBA (Supplementary Figures 3 and 4). Supplementary Table 2 lists the association p-values between genes and RBA (DOI:10.5281/zenodo.3786854).

**Fig. 1.**
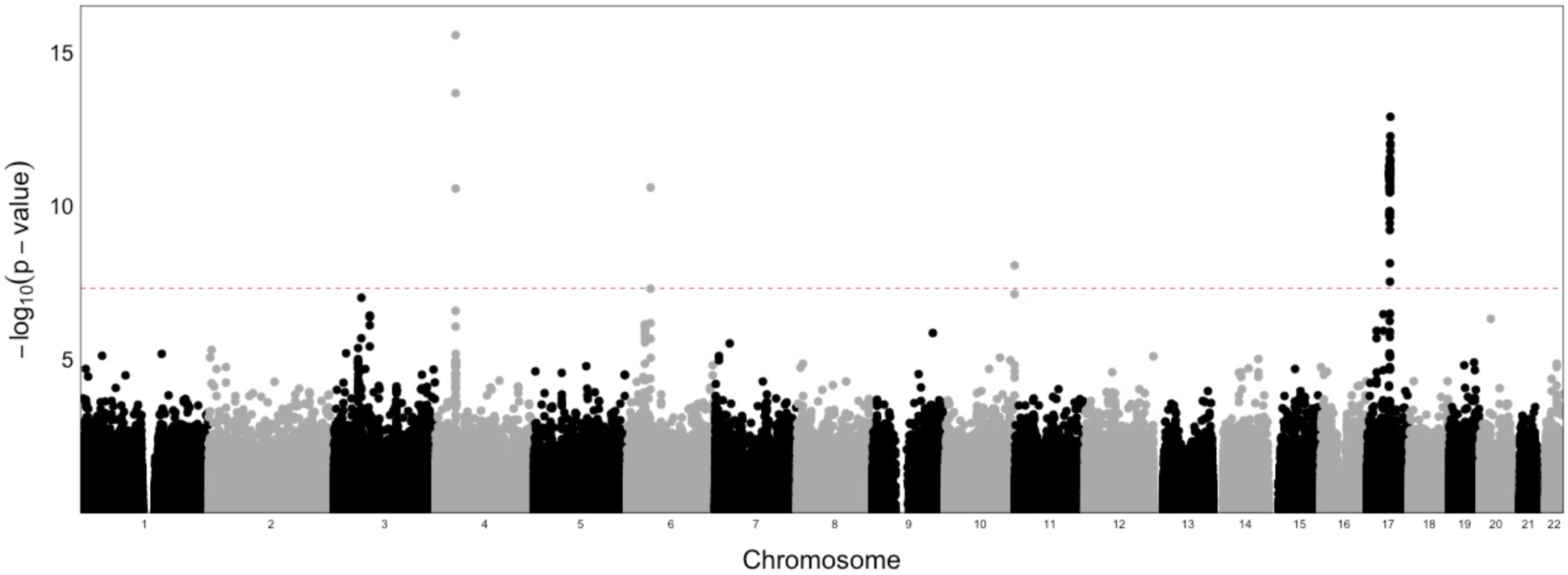
Manhattan plot of the association p-values between SNPs and relative brain age across the genome. The red dashed line indicates the genome-wide significance threshold on p-value (i.e., 5E-8).

**Fig. 2.**
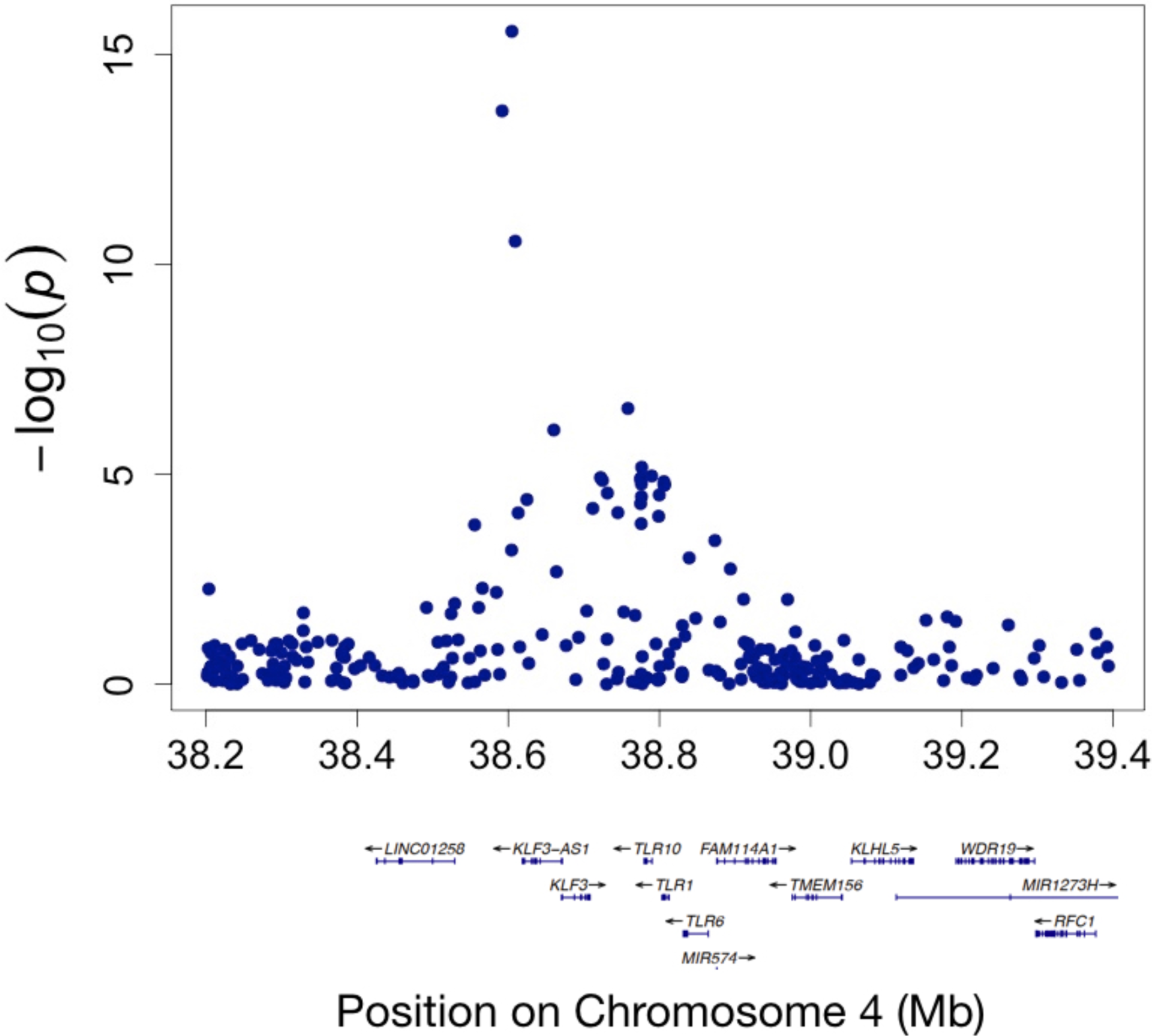
Regional visualization of a locus on Chromosome 4 where SNPs showing genome-wide significant associations with relative brain age are located.

### 2.3 Comparing CNN and linear regression models

The CNN model had a significantly smaller MAE between PBA and chronological age than the linear regression model, whose median MAE was 3.7 years in the five evaluation sets (paired t-test p-value < 0.05). The SNP-RBA association p-values were stronger when using RBA derived from the CNN model than using RBA from the linear regression model (Fig. 4). Also, when using RBA from the linear regression model, only the SNPs from a chromosome 17 locus were significantly associated with RBA (Supplementary Fig. 2).

**Fig. 3.**
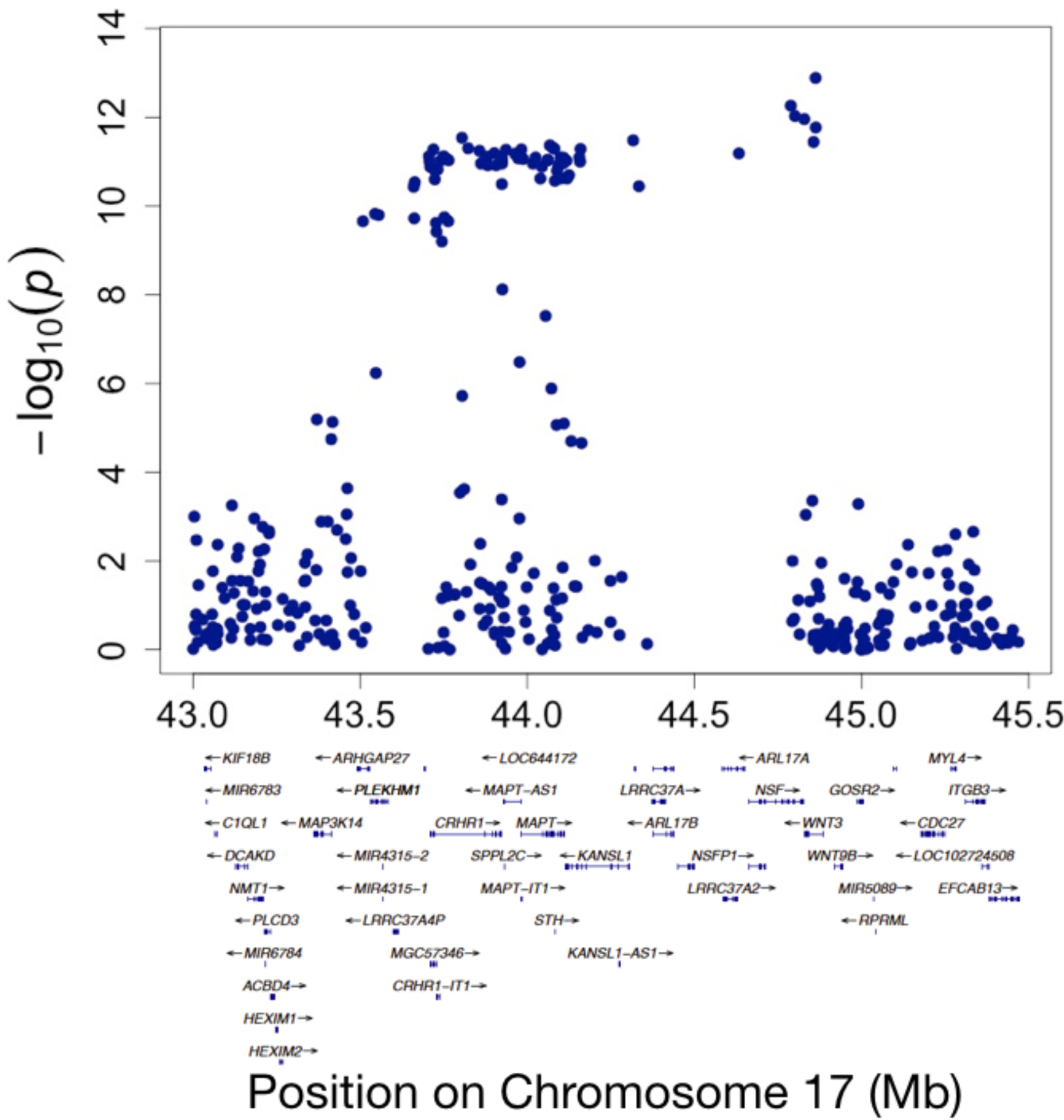
Regional visualization of a locus on Chromosome 17 where SNPs showing genome-wide significant associations with relative brain age are located.

**Fig. 4.**
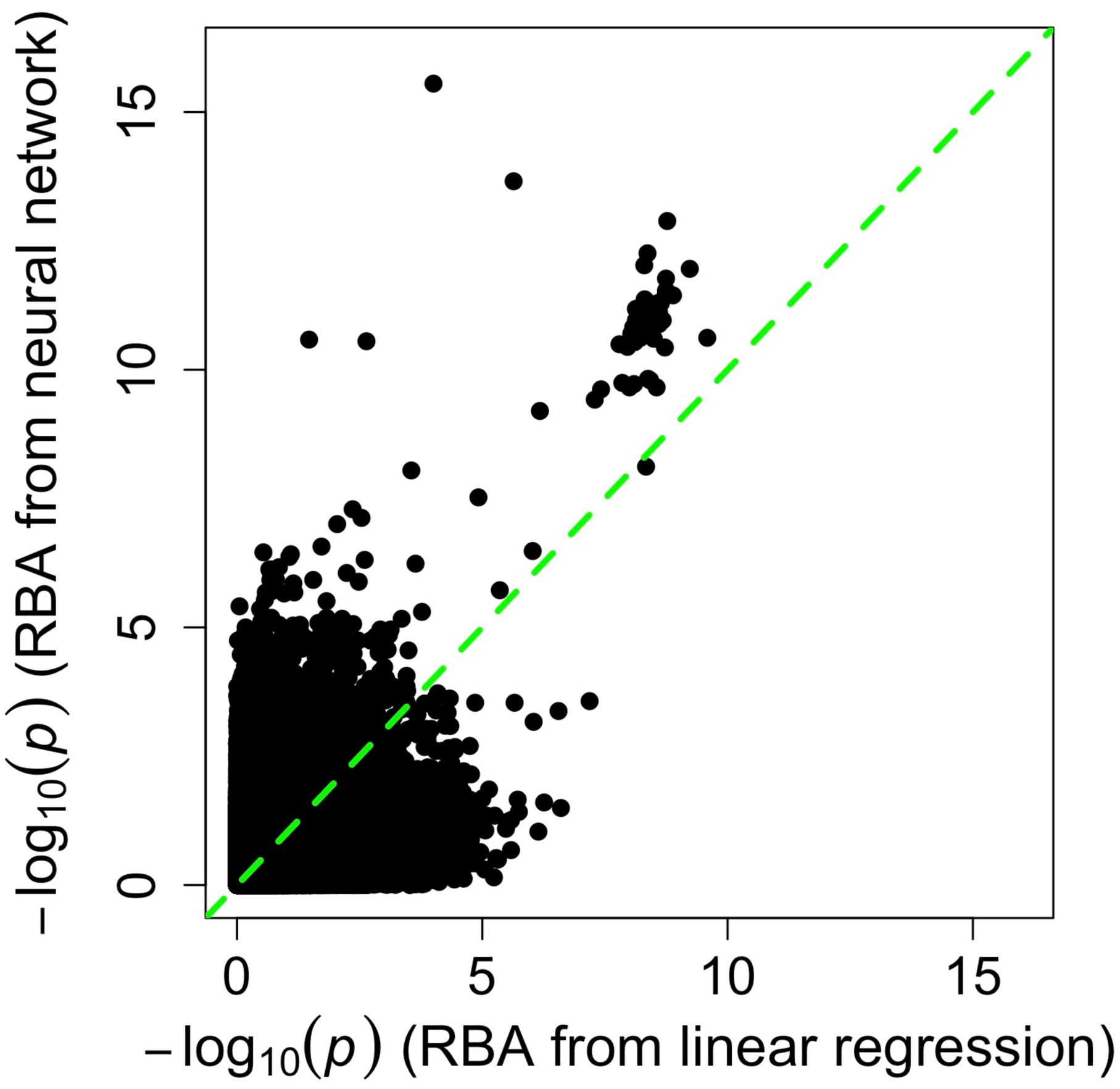
Comparison of SNP-RBA association signals identified using RBA from a linear regression model and using RBA from a convolutional neural network model.

## 3 Discussion

In this study, we trained a CNN for obtaining PBA and further studied associations between SNPs and brain aging. The median MAE between PBA and chronological age was 2.7 years in the evaluation sets. The metric representing the difference between PBA and CA (PBA-CA) was negatively associated with chronological age when using CNN model (Supplementary Fig. 1). This negative association was similar to the pattern found between PBA-CA and chronological age where PBA was developed using a linear regression model (*5, 6*). This negative association is likely a result of regression dilution and has been discussed previously by Smith et al. (*6*) Deriving PBA using a CNN does not eliminate this issue, since the last layer of the CNN model was fundamentally a linear regression model. Therefore, to adjust for the undesirable association between (PBA-CA) and chronological age, we further derived RBA after obtaining PBA and assessed association between RBA and SNPs.

We found SNPs from four loci to be significantly associated with RBA. The SNP most significantly associated with RBA (rs337638, p-value = 3E-16) was located in a non-coding region on chromosome 4. The protein coding gene located the closest to this SNP is KLF3. KLF3 mediates lifespan in C. elegans (*12*) and is possibly associated with educational attainment in human subjects (*13*). SNPs in a chromosome 17 locus also showed significant associations with RBA. In this locus, genes including MAPT, NSF, and WNT3 have significant gene-level associations with RBA and are of particular interest (Supplementary Table 2). Previous studies have shown that mutations in MAPT, which encodes tau protein, are associated with dementia and Parkinson’s disease (*8*). The NSF gene encodes ATPase that is involved in cellular membrane fusion events, and is associated with neuronal intranuclear inclusion disease (*14*). The WNT3 gene is associated with general cognitive funcrtion (*15*). In addition, the SNP rs2677110 located in the RUNX2 gene on chromosome 6 and the SNP rs12775208 located in the NKX6-2 gene on chromosome 10 also showed significant association with RBA. RUNX2 gene encodes a transcription factor that regulates the expression of osteocalcin (*16*). Interestingly, low osteocalcin level is associated with age-related decline in cognitive function, which may contribute to the association between this RUNX2 variant and RBA (*17, 18*). NKX6-2 encodes a transcription factor that regulates myelin sheath formation and maintenance in the central nervous system (*19, 20*). While there are plausible links between these variants and overall brain health, the biological functions of these RBA-associated SNPs remain to be further investigated to understand how they affect brain aging.

We further compared the results based on CNN-derived PBA to those based on regression model-derived PBA. In our analyses, the CNN model was more accurate than a linear regression model in predicting brain age (median MAE 2.7 years versus 3.7 years). The SNP-RBA association signals were stronger (i.e., with more significant p-value) when using RBA derived from the CNN model than using RBA derived from the linear regression model (Fig. 1, Fig. 4, and Supplementary Fig. 2). Nevertheless, the additional cost of time to train a CNN model was not trivial. For our training process, the CNN model took about two days to converge (with two NVIDIA GPUs). As a comparison, training a linear regression model with FreeSurfer measurements as predictors for brain age only took two minutes on one CPU.

Besides genetic factors, lifestyle habits and diseases are also associated with brain aging. For example, heavy smoking and alcohol consumption are associated with accelerated brain aging in both specific brain regions and overall structural brain aging (*5*). Physical exercise improves general health and may also slow down brain aging and reduce Alzheimer’s disease risk (*21, 22*). Diseases such as diabetes and Schizophrenia are associated with accelerated brain aging (*7, 23*). As large-scale data including both brain imaging data and information of these factors become accessible, RBA based on NN models can be used to better characterize the links between these factors and brain aging.

In summary, the CNN model more accurately estimated PBA and RBA than a regression model, which lead to identification of novel associations between SNPs and brain aging. A more accurate RBA is likely to lead to better characterization of the association between brain aging and other lifestyle or disease factors.

## 4 Material and methods

### 4.1 Summary of samples used

A sample of 16,998 subjects from the UK Biobank study with European ancestry who had both T1 MRI brain scans and genetic data were included in the study. Subjects’ ages ranged from 45 to 81 years (mean = 63 years), with 53% identified as female. The brain scans were registered to MNI 152 space (*24*). The genetic data included 538,476 quality controlled SNPs. Details of quality control on brain imaging data and genetic data has been described previously (*5, 25*).

### 4.2 Obtaining predicted brain age and relative brain age based on a CNN model

#### 4.2.1 Training CNN models for predicting brain age

We used a five-fold cross-validation strategy for training a CNN model for predicting brain age and for doing SNP-RBA association analyses. Specifically, we randomly split the subjects into five batches of 20% of the data, with each having the same distribution of age and gender. We used 80% of the samples (i.e., four batches) for training a CNN model for predicting brain age (PBA), and then applied the trained models to the remaining 20% of samples and obtained PBA of these samples. We crossed over the split samples and obtained PBA for each batch of the 20% of subjects.

#### 4.2.2 Details of the CNN model

We used a CNN model with ResNet structure (*26*) implemented in NiftyNet for predicting brain age based on 3D MRI data (https://niftynet.io). We chose the ResNet structure because empirical evidence has shown that it is easy to optimize and performs well in imaging data analyses (*26*). We down-sampled the MRI data from a resolution of 182×218×182 to 91×109×91 before training the CNN, due to the GPU memory limitations. The ResNet model provided by Niftynet comprised of a convolution layer, six bottleneck layers, a fully connected layer, and an output layer. Multiple short-cut connections were added among bottleneck layers. We specified the structure as follows. The initial convolution layer contained 64 filters. There were six bottleneck layers: bottleneck layer 1 and bottleneck layer 2 had 128 filters; bottleneck layer 3 and bottleneck layer 4 had 256 filters; bottleneck layer 5 and bottleneck layer 6 had 512 filters. Random rotations between -5 and 5 degrees were included as a data augmentation strategy. The model was trained with learning rate at 0.0001 on two GPUs. The CNN structure is illustrated in Fig. 5.

**Fig. 5.**
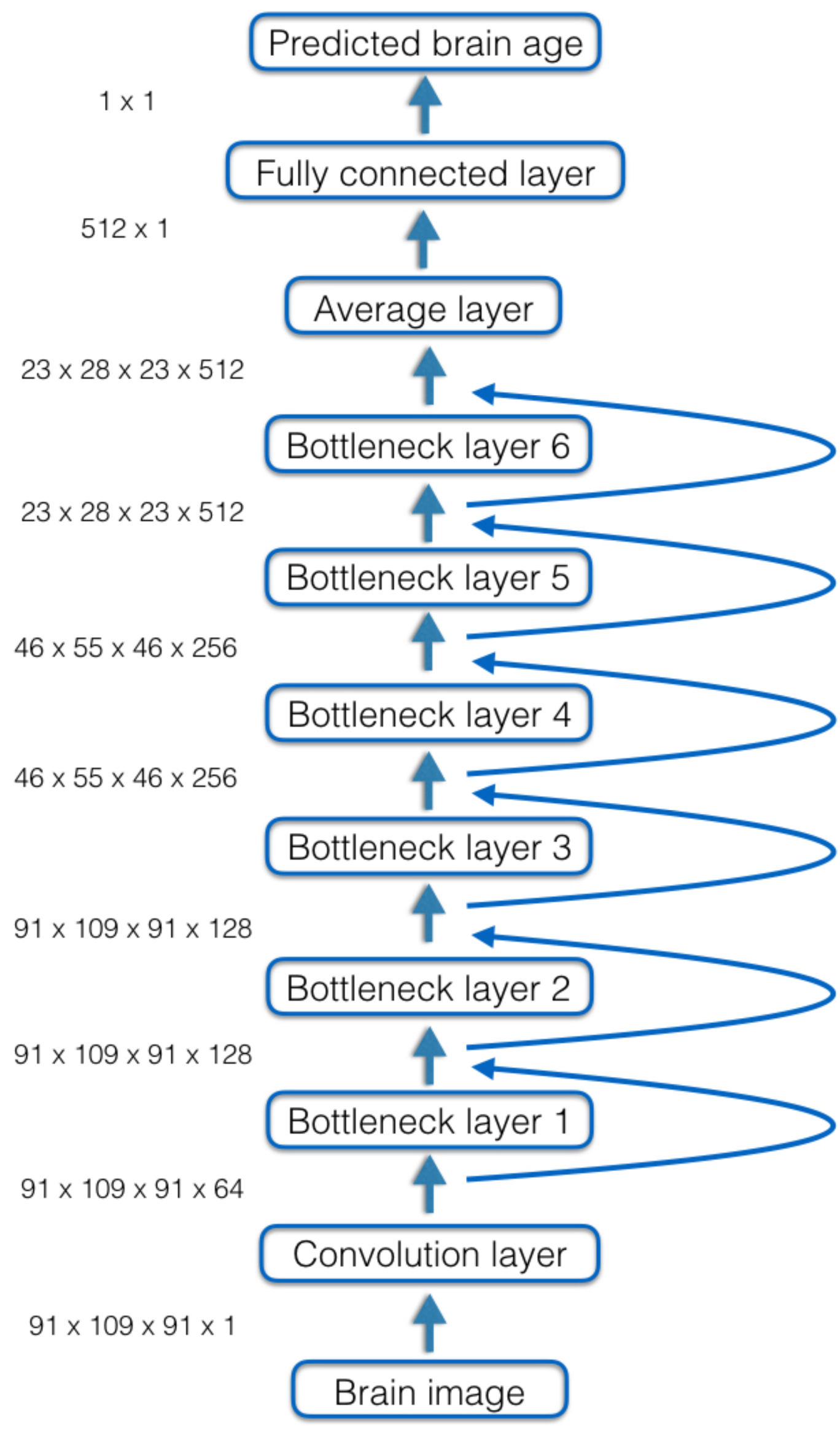
Structure of the convolutional neural network model for predicting brain age based on MRI data. Curved arrows indicate skip connections between layers.

#### 4.2.3 Obtaining relative brain age

The metric representing the difference between PBA and CA (PBA-CA) was negatively associated with chronological age when using CNN model (Supplementary Fig. 1). Therefore, we further derived RBA after obtaining PBA (as described by Ning et al. (*5*)). We then combined RBA of all evaluation set subjects and assessed association between RBA and SNPs. Fig. 6 illustrates the procedure for obtaining PBA and RBA through a 5-fold cross-validation strategy and then carrying out RBA-SNP association analyses.

**Fig. 6.**
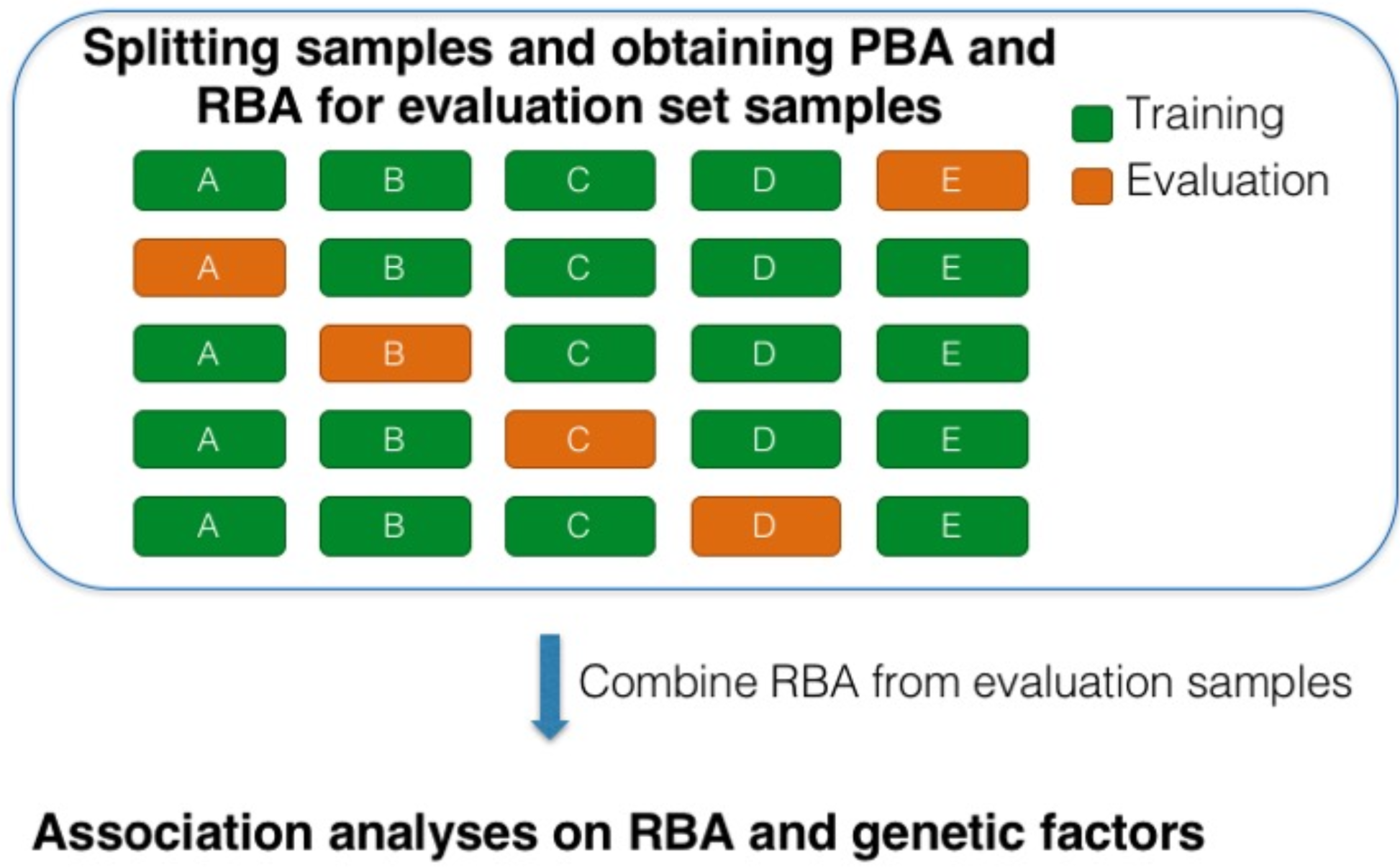
Procedure for obtaining PBA and RBA through a 5-fold cross-validation strategy and then carrying out RBA-SNP association analyses.

### 4.3 Obtaining predicted brain age and relative brain age based on a linear regression model

We trained linear regression models with LASSO regularization to obtain PBA using the same training and evaluation sets as used for the CNN models. In the linear regression models, the predictors were 403 brain morphometric measurements from Freesurfer (*27*) (e.g., volume of cortical, subcortical, white matter, and ventricle regions, thickness and surface area of cortical regions, intracranial volume, etc.), and the response variable was CA. After obtaining PBA, we further derived RBA metric. Details of obtaining PBA and RBA using the regression model are described in a previous paper (*5*).

### 4.4 SNP-RBA association analyses

We used a linear mixed model implemented in BOLT-LMM (*28*) to test the association between SNPs and RBA, adjusting for gender, the first three genetic principal components, and batch information during sampling. We further used FUMA (*29*) to obtain gene-level association with RBA.

## Supplementary Figures

**Supplementary Fig. 1.**
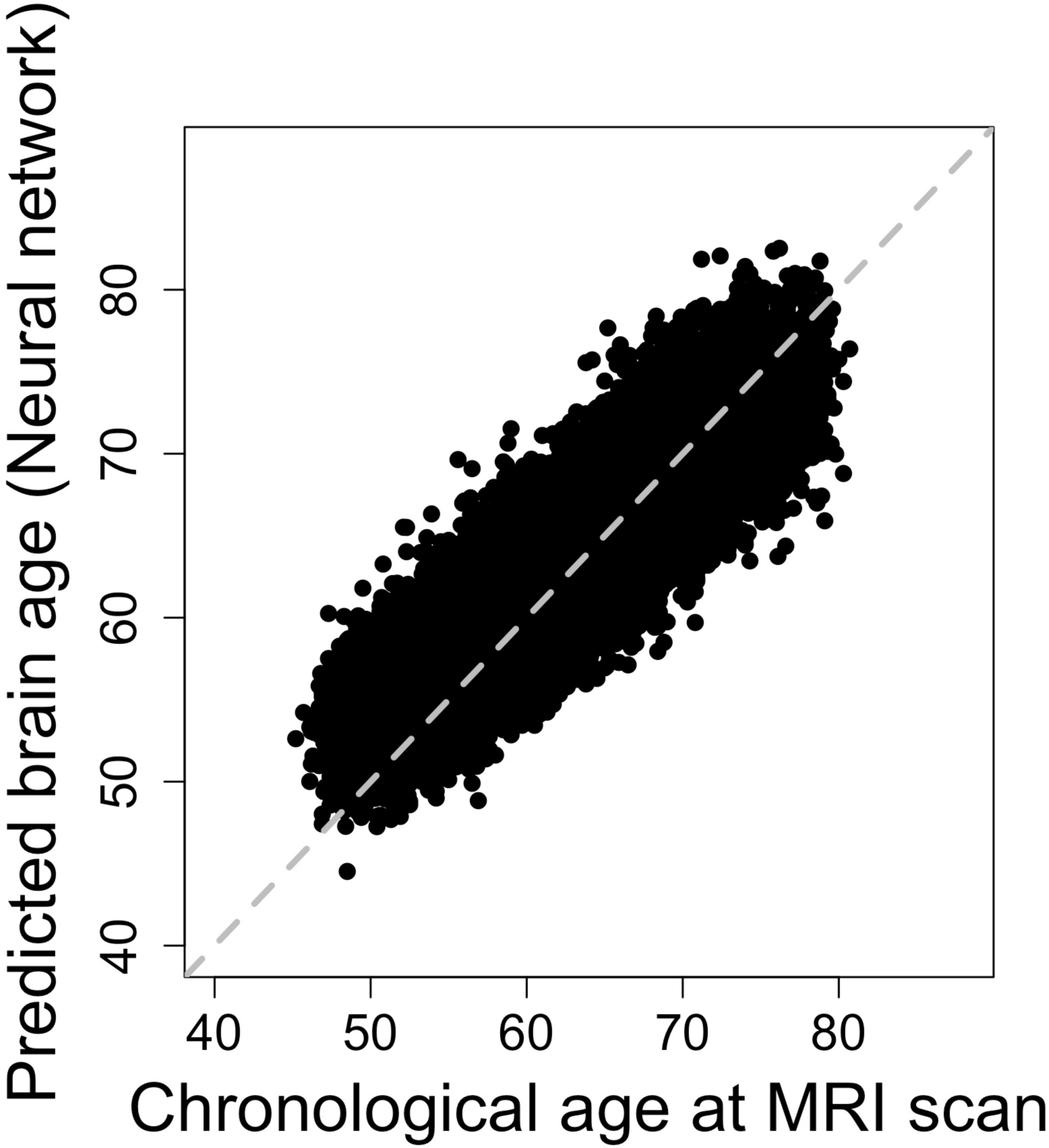
Association between chronological age and predicted brain age from a convolutional neural network model.

**Supplementary Fig. 2.**
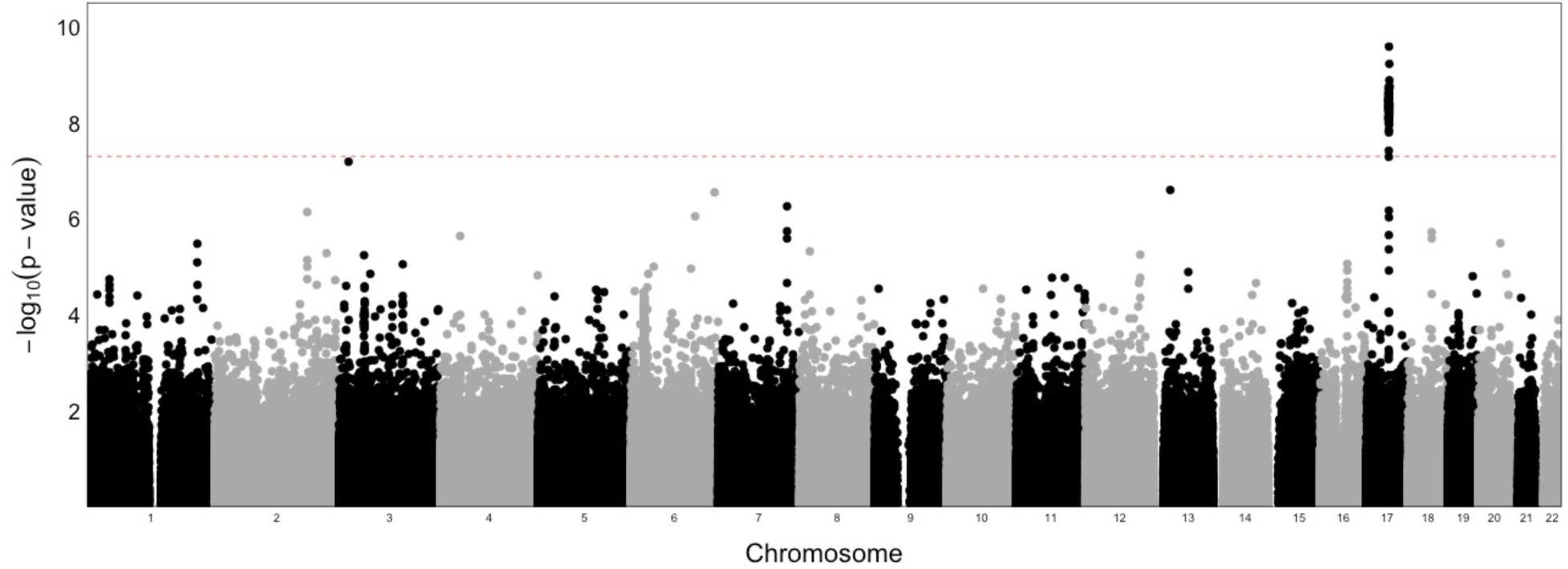
Manhattan plot of the association p-values between SNPs and relative brain age across the genome, where RBA is derived from a linear regression model. The red dashed line indicates the genome-wide significance threshold on p-value (i.e., 5E-8).

**Supplementary Fig. 3.**
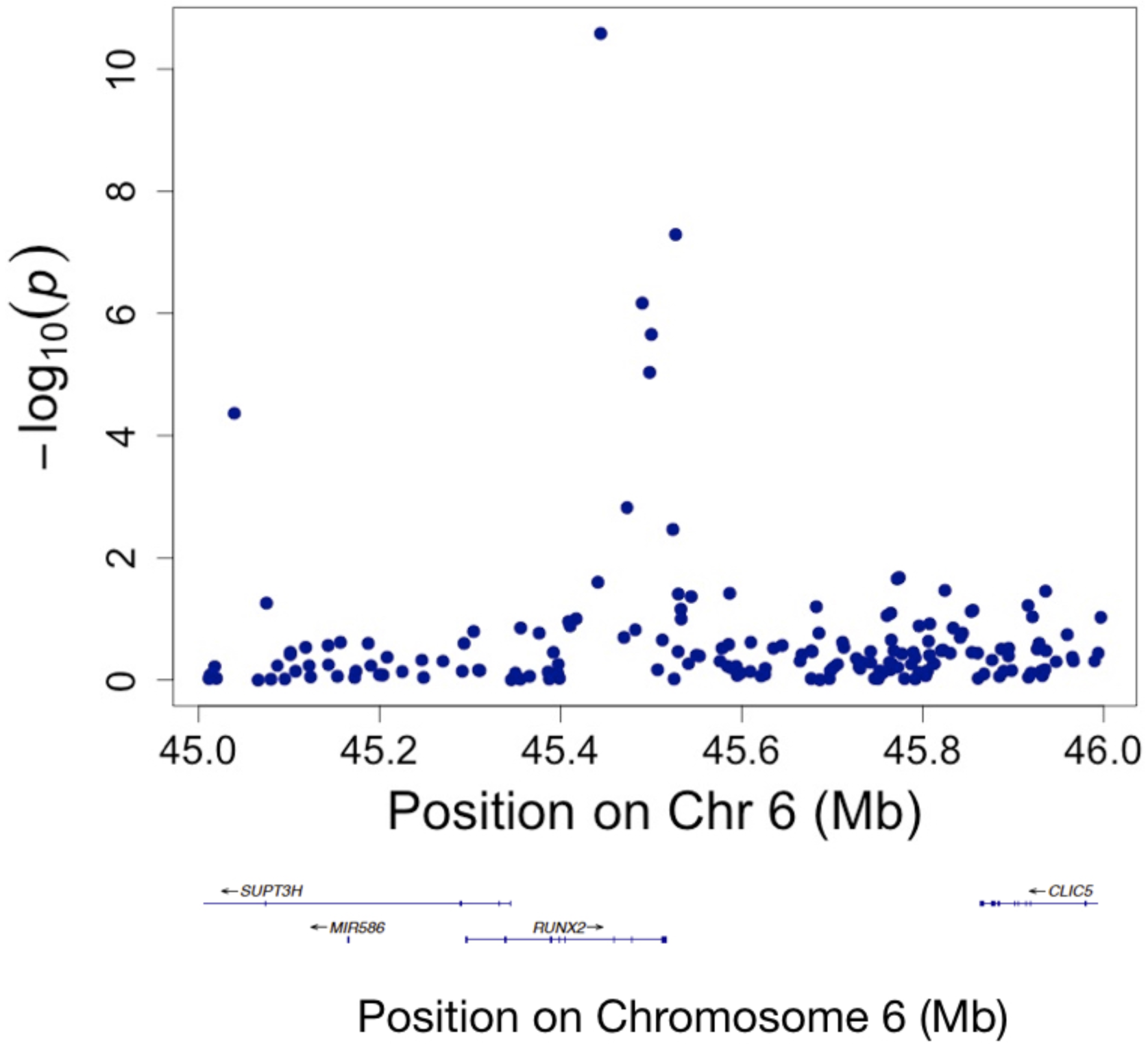
Regional visualization of a locus on chromosome 6 where a SNP showing genome-wide significant associations with relative brain age is located.

**Supplementary Fig. 4.**
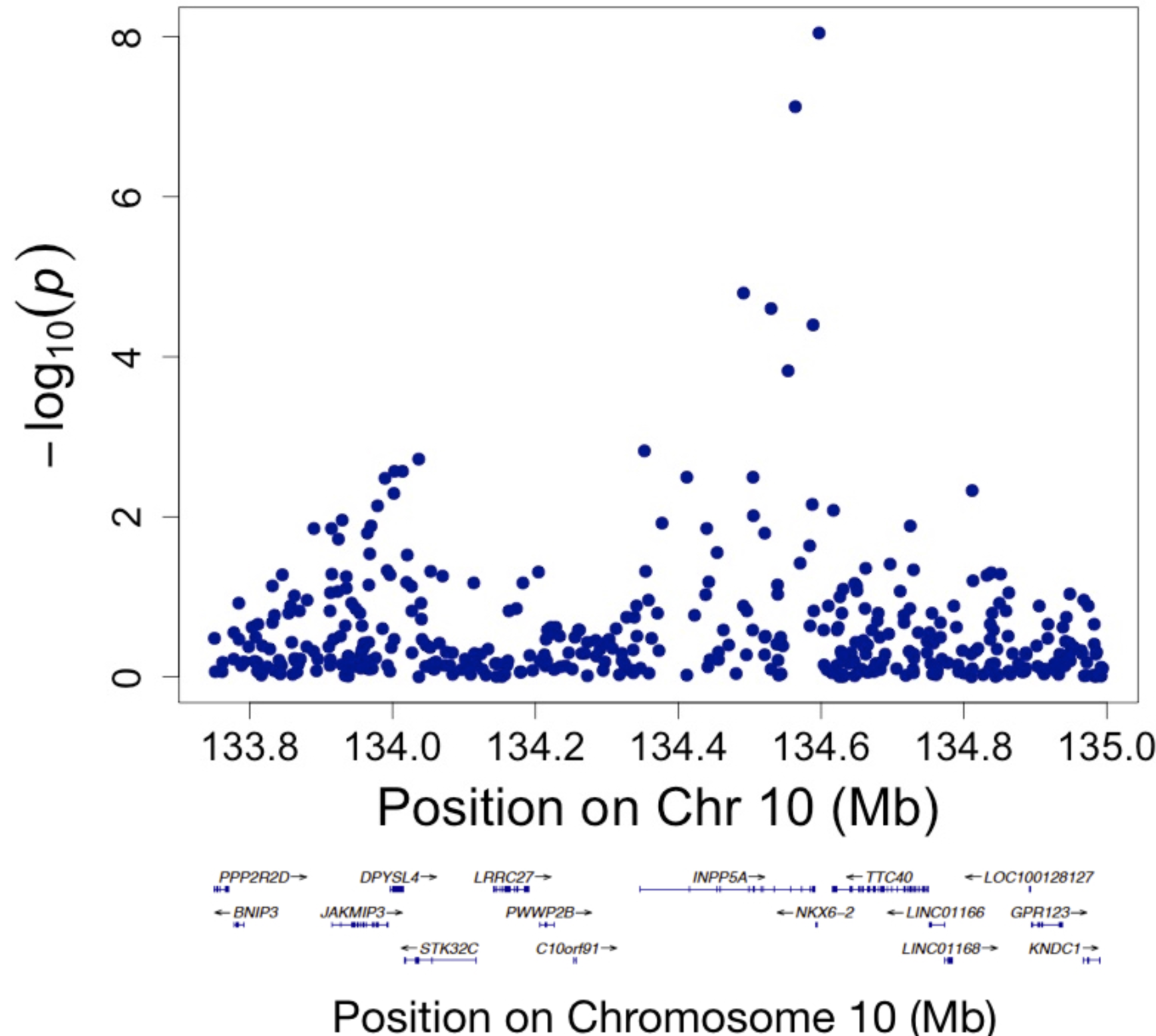
Regional visualization of a locus on chromosome 10 where a SNP showing genome-wide significant association with relative brain age is located.

## Acknowledgements

We thank Dr. Bo Chen for helpful discussions on data analyses procedure. We also acknowledge the contributions of members of the UK Biobank coordinating center.

